# Differential Regulation Analysis Quantifies Mirna Regulatory Roles and Context-Specific Targets

**DOI:** 10.1101/2022.07.24.501303

**Authors:** Boting Ning, Tamar Spira, Jennifer E. Beane, Marc E. Lenburg

## Abstract

Rewiring of transcriptional regulatory networks has been implicated in many biological and pathological processes. However, most current methods for detecting rewiring events (differential network connectivity) are not optimized for miRNA-mediated gene regulation and fail to systematically examine predicted target genes in study designs with multiple experimental or phenotypic groups. We developed a novel method to address these shortcomings. The method first estimates miRNA-gene expression correlations with Spatial Quantile Normalization to remove the mean-correlation relationship. Then, for each miRNA, genes are ranked by their correlation strength per experimental group. Enrichment patterns of predicted target genes are compared using the Anderson-Darling test and significance levels are estimated via permutation. Finally, context-specific target genes for each miRNA are identified with target prioritization based on the correlation strength between miRNA and predicted target genes within each group. In miR-155 KO RNA-seq data from four mice immune cell types, our method captures the known cell-specific regulatory differences of miR-155, and prioritized targets are involved in functional pathways with cell-type specificity. Moreover, in TCGA BRCA data, our method identified subtype-specific targets that were uniquely altered by miRNA perturbations in cell lines of the same subtype. Our work provides a new approach to characterize miRNA-mediated gene regulatory network rewiring across multiple groups from transcriptomic profiles. The method may offer novel insights into cell-type and cancer subtype-specific miRNA regulatory roles.

## Introduction

Gene expression regulation within cells can be modeled via networks, where the transcriptional regulators, such as transcription factors (TFs), and their downstream target genes are represented as nodes and regulatory relationships are represented as edges^1,2^. Through tight control of the transcriptional network, cells can accurately coordinate gene expression and establish correct cellular functions under normal conditions. In the meantime, the gain or loss of connectivity in the transcriptional network, or “rewiring” events, can result in altered gene expression profiles observed among differentiating or treated/perturbed cells or between cancer-subtypes^3–6^. Computational methods have been developed to detect and statistically quantify transcriptional network rewiring events between two groups from bulk gene expression profiles. These methods utilize gene-gene expression correlation coefficients, graphical models, or Latent Dirichlet allocation with TF chromatin-binding profiles to infer the transcriptional regulator (typically TFs) whose connectivity with downstream targets is significantly rewired^7–10^.

MicroRNA (miRNA) is a class of short, non-coding RNAs which utilizes complementary sequence paring between its seed sequence and the 3’ untranslated region (UTR) of gene transcripts to repress target gene expression levels^11–13^. Through acting as post-transcriptional regulators, miRNAs participate in a wide range of biological and cellular processes, including cell differentiation, development, and carcinogenesis processes^14–17^. It has been shown that miRNA may undergo network rewiring and have different functions between cell-types or cancer molecular subtypes^18–21^. Particularly shown by the study from Hsin *et al*., the rewiring of miRNA regulatory networks between cell-types is predominantly due to miRNA target binding switching, rather than 3’UTR isoform or expression levels^22^. While these studies provided valuable knowledge on miRNA regulatory networks, they often rely on single-cell RNA sequencing or complicated functional profiling such as CLIP-seq with Halo-enhanced Ago2 pull-down^23^ or CLASH^24^, which is expensive and not always feasible. Thus, it is advantageous to develop methods capable of identifying miRNAs with differential regulatory roles across either multiple cell-types or cancer subtypes from bulk miRNA and gene expression profiles.

Most current computational methods for detecting rewiring events, are not optimized for miRNA-mediated gene regulation and cannot be directly applied to scenarios involving miRNAs for several reasons. 1) Previous methods were developed to build gene-gene networks (all nodes connected and associations between nodes could be either positive or negative), but did not include the miRNA predicted target information nor the miRNA gene repression activity in the computational model; 2) Previous methods outputted either single miRNA-gene connection or single module consisted of multiple interconnected miRNAs/genes rather than potential miRNAs; 3) Previous methods focus on comparisons between two groups, whereas when comparing between cell-types or cancer molecular subtypes, researchers are often dealing with more than two groups.

Here, we present a novel computational framework and R package *<u>Di</u>fferential <u>Re</u>gulation <u>A</u>nalysis of <u>mi</u>RNA* (DReAmiR; https://github.com/ningb/DReAmiR) to address the aforementioned challenges. Integrating mRNA and miRNA expression profiles with the predicted target information, DReAmiR first estimates miRNA-mRNA correlation matrices per group while removing the bias from the mean-correlation relationship^25^. Then, it identifies miRNA with significantly context-specific targets across multiple experimental groups based on differential enrichment of predicted targets between groups using genes ranked by mRNA / miRNA correlation coefficients. Finally, DReAmiR can prioritize target genes uniquely regulated in each group using either a graph embedding-based approach or iteratively maximizing the difference in enrichment score (dES).

## Methods

### DReAmiR

The DReAmiR workflow is divided into three main steps (**Figure 1**): (1) Remove mean-correlation relationship using spatial quantile normalization (SpQN), (2) Identify miRNAs with significant context-specific targets with differential regulation analysis, and (3) Prioritize the target genes specifically regulated by a miRNA in each group.

**Fig 1.**
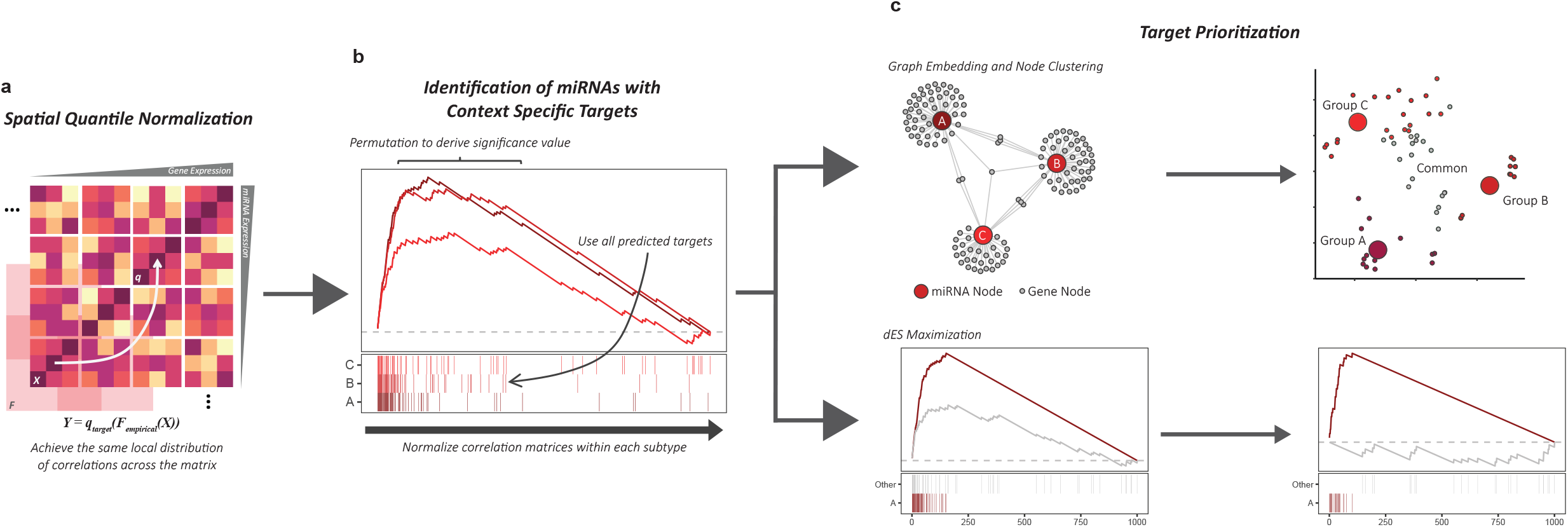
DReAmiR method overview. DReAmiR conisists of three major steps. **a**. SpQN removes the mean-correlation relationship in the miRNA-gene correlation matrices. **b**. Differential regulation analysis identifies miRNAs with context-specific target genes by comparing the ranking of miRNA-gene correlation coefficients within each group. **c**. Two methods were developed to prioritize experimental group-specific target genes.

#### SpQN

To remove the bias from gene/miRNA expression levels on correlation estimations (the mean-correlation relationship) in the miRNA-gene correlation matrix, we modified the SpQN algorithm developed by Wang Y. *et al*. (2020) to accommodate an asymmetrical matrix^25^. Briefly, a Pearson correlation matrix is first constructed for each experimental group between miRNAs and genes and sorted by miRNA and gene expression levels. Then, the correlation matrix is separated into non-overlapped sub-matrices based on gene and miRNA expression levels. Next, for each non-overlapped sub-matrix, a larger overlapping matrix is used for estimating the empirical correlation coefficient distribution. The number of bins for genes and miRNAs and the overlap size can be specified by the user. By default, DReAmiR separates the correlation matrix into 20×20 submatrices and constructs overlapping matrices with 1000 genes and 150 miRNAs each, which are suitable for typical RNA and small-RNA sequencing experiments in which about 10-20 thousand genes and around one thousand miRNAs can be detected. These parameters are chosen to balance the smoothness in normalization and the total running time. Using the sub-matrix close to the top right corner as reference (by default, the sub-matrix corresponding the second-highest miRNA and gene expression), 1-dimensional quantile normalization is applied to match the correlation density distribution of all other bins. Further information on parameter tunings for SpQN can be found in Wang Y. *et al*. (2020)^25^.

#### Differential regulation analysis

After correcting for the mean-correlation relationship, differential regulation analysis can be performed to identify miRNAs with significant context-specific target genes. This step requires two inputs: the correlation matrices for each group and the predicted target genes of miRNAs. For each miRNA, the genes are first ranked by their normalized correlation coefficients to the miRNA, from negative to positive, to generate a gene rank list for each group. Then, the enrichment pattern of predicted target genes in each rank list is compared using the Anderson-Darling (AD) test. Permutation of group label is performed and the observed test statistic is compared to the distribution from permutations to generate empirical p-values. When only two groups are present, Kolmogorov-Smirnov (KS) test is used instead. Post-hoc analysis using the KS test can be performed afterward to examine pair-wise significance.

#### Target Prioritization

For the miRNAs whose regulated genes are significantly different between groups, DReAmiR can further prioritize their target genes in the leading edge of each group based on association strength, assuming that being in the leading edge suggests a gene is strongly regulated by the miRNA in at least one group. DReAmiR provides two methods for this purpose.

1. The graph embedding and node clustering method aims to classify and label the target gene clusters as either group-specific or shared between groups. First, a network is constructed with nodes representing genes and group-specific miRNAs. An edge between a gene node and a group-specific miRNA node represents the gene being in the leading-edge of that group based on differential regulation analysis, with the association strength as the edge weight. Then, the large information network embedding (LINE) algorithm^26^ is used to identify embedded features. We chose LINE over other network embedding methods due to its scalability and ability to preserve both local and global network structures. Fuzzy k-means clustering is performed within the embedding space to cluster the target genes. Clusters containing one or more group-specific miRNA nodes are labeled accordingly. For each of the remaining clusters, a one-tail KS test is performed to compare whether the node membership density associated with each of the labeled clusters is significantly higher than the other labeled clusters, and the cluster labels are then assigned based on the number of significant results. A cluster with membership density significantly higher for one labeled group is labeled as a uniquely regulated target cluster for that group. A cluster with more than one significant result is labeled as a shared target cluster. Cluster without any significant result is a common target gene cluster regulated equally across all groups.
2. Difference in enrichment score (*dES*) maximization method aims to find the set of target genes that are most strongly regulated in the group of interests compared to the reference group. The reference group can either be another group that the user wants to compare to, or all other groups combined. Starting with all the genes in the leading edge in the group of interests (notated as 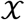) as the baseline target set, DReAmiR first calculates the *ES_normalized_*, which is the observed enrichment score divided by the mean of permutation *ES* values from shuffling the rank list 500 times. Then, *dES* is derived as 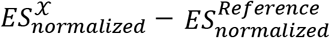. Then, each predicted target gene in the baseline target set is removed one at a time and the changes in *dES* are calculated. The gene that yields the largest increase in *dES* is then removed to form the new target set. The process is repeated until no gene remains in the target set or the minimum target size specified by the user is reached (by default 20). The *dES* calculated at each step is recorded and back-traced. The set of genes giving the largest *dES* is selected as the final target set for the group of interest, which yields the largest separation in enrichment score compared to the reference group. Of note, the direction of enrichment to test, either positive or negative, can be selected by the user and the calculation in *ES* calculation is changed accordingly.

### Other Functions

To facilitate the workflow, DReAmiR also contains several utility functions:

#### Correlation matrix and predicted target gene matrix construction

To simplify the workflow and ensure the reproducibility of the analysis, DReAmiR only requires three simple inputs: gene expression matrix, miRNA expression matrix, and the group label. DReAmiR examines the format of these inputs and returns group-specific correlation matrices for the samples with complete records. In addition, with the help of multimiR Bioconductor package^27^, users can specify which and the number of miRNA target databases to use. The intersection of databases will be used for miRNA-gene target information. For example, if a gene is recorded as the target of a miRNA by 3 out of the 5 miRNA target databases, it will be retained for down-stream analysis. DReAmiR directly generates a binary target matrix for miRNA-gene pair with the same size as the miRNA-gene correlation matrix.

#### Visualization

The output from the differential regulation analysis can be visualized as a multi-group enrichment plot (**Fig 2a** and **Fig 3b**). The plot contains two panels: the top panel depicts the enrichment patterns and the enrichment score per group, and the bottom panel depicts the position of predicted target genes in each rank list.

**Fig 2.**
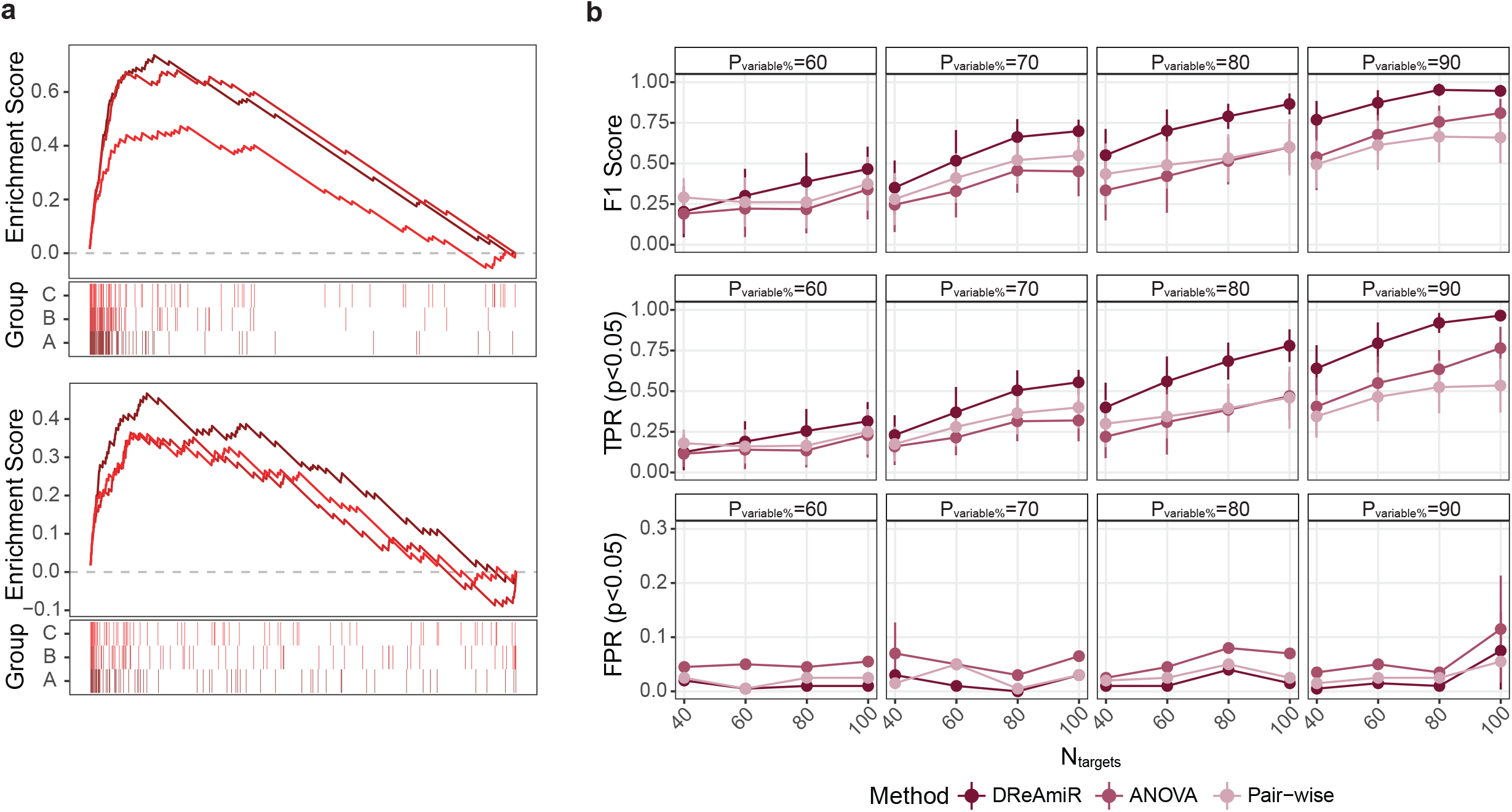
DReAmiR performance in simulated data. **a.** Enrichment plots for simulated true positive (top) and true negative (bottom) differentially regulating miRNAs. **b.** Performance comparisons between DReAmiR, ANOVA and pair-wise comparison in the simulated data with sample size per group equaled 70. The simulation was performed across different simulation parameters, including the number of predicted target genes per miRNA, and the percentage of variable targets per group. F1 score (F1), true positive rates (TPR), and false positive rates (FPR) across 20 iterations per combination of simulated parameters were evaluated by the mean and standard deviation.

**Fig 3.**
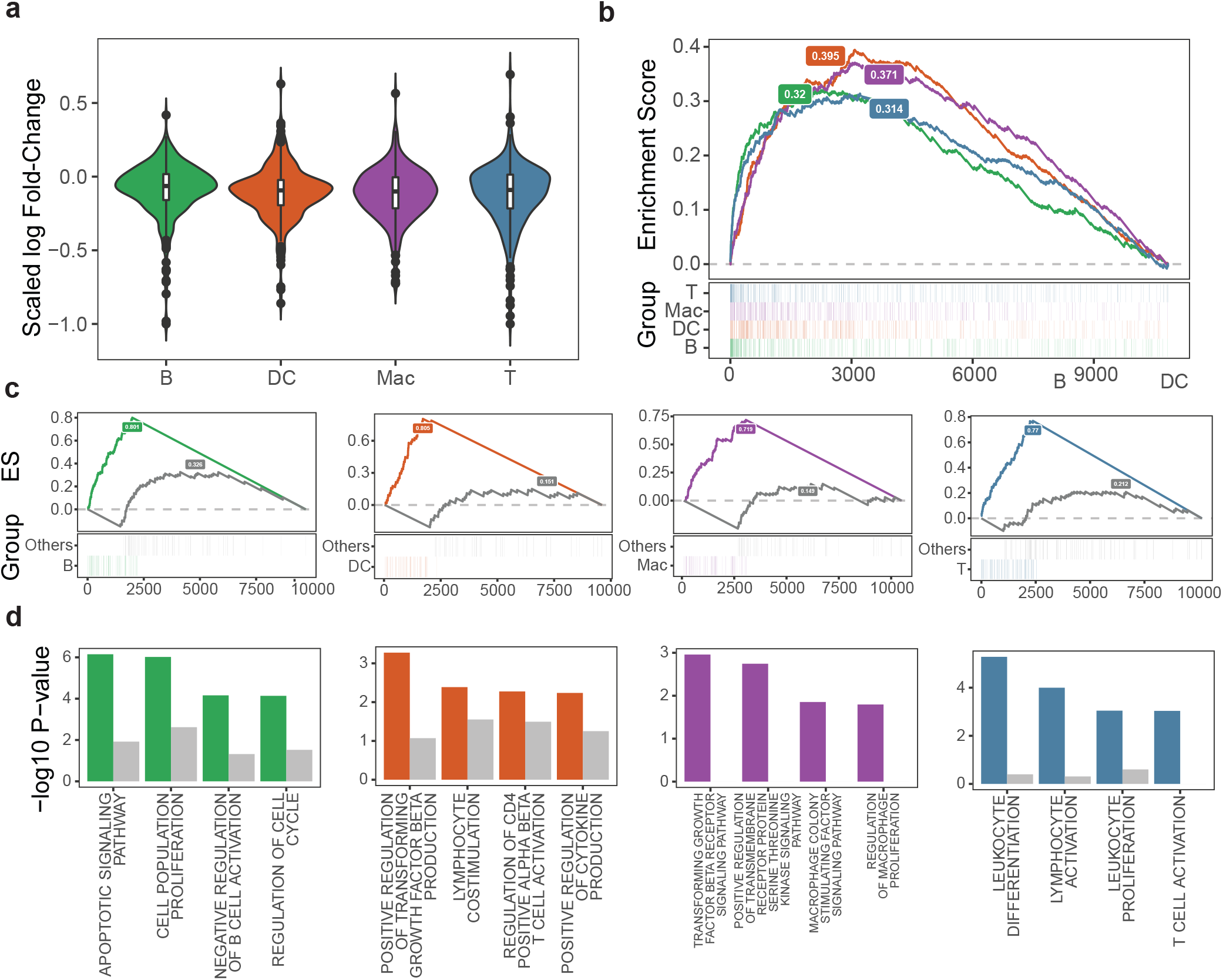
DReAmiR identifies miR-155 target genes involved in functional pathways with cell-type specificity. **a.** Enrichment plot of miR-155 predicted target genes in the gene rank list sorted by logFC in miR-155 KO samples comparing to the controls within each immune cell-types (Permutation p-value = 0.01). **b.** logFC densities of miR-155 predicted target genes in miR-155 KO samples compared to the controls within each immune cell-types (ANOVA p-value = 0.08). **c.** Enrichment plots for the miR-155 prioritized target genes per immune cell-type, using the max-dES method. For each cell-type, the gene list ranked by logFC associated with miR-155 in the other three cell-types compared to the were used as the reference group. **d.** Functional pathway enrichment results for the prioritized miR-155 target genes per cell-type (colored), and the differentially expressed miR-155 target genes following miR-155 KO (grey).

#### Parallelization

DReAmiR contains several computationally intensive tasks, including the group-wise correlation matrix calculation, the SpQN, and max-dES with iterative searches. To reduce the analysis time, we implemented parallelization for these steps when users have access to multi-core computation.

### Data Simulation

To evaluate the performance of DReAmiR in detecting miRNAs with different regulatory roles across multiple groups, we generated sample matched gene and miRNA expression where the predicted target genes of a miRNA have different patterns of negative enrichment along the gene rank list by miRNA-gene correlation strength. The simulation dataset contained three groups of samples with expression profiles of 1000 genes and 20 miRNAs. 10 miRNAs were assigned to have context-specific target genes between groups (true positive) and 10 to be not different (true negative). For each miRNA, we first randomly picked *N_target_* genes to be the predicted targets, mimicking the information from miRNA predicted databases. The covariance between the predicted target genes and miRNAs was simulated by a normal distribution with negative mean 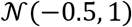 representing globally suppressive effects on gene expression by miRNAs. Then, for those miRNAs assigned to be true positive, we randomly picked *P_variable%_* percent of the predicted targets per group to be the group-specific target and should be negatively regulated more strongly within that group. The association between group-specific target genes and miRNA was simulated by shifting the covariance towards the negative direction proportionally from the mean within two groups with a left-skewed beta distribution Beta(*α*, *β*) where *α* = 20 and 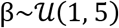. Next, the nearest positive-definite of the covariance matrix is calculated using the ‘make.positive.definite’ function from lqmm R packages^28^. The gene and miRNA expression matrices were then simulated for *N_sample_* per group by multivariate Gaussian distribution 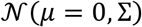 using ‘mvrnorm’ function form the MASS R package^29^. Finally, the miRNA-gene correlation matrix was calculated and exported along with the predicted targets per miRNA as the input for performance evaluation.

Evaluation of DReAmiR performance was conducted by simulating datasets with the following parameters:

*N_target_* = {40, 60, 80, *and* 100}
*P_variable%_* = {60, 70, 80, *and* 90}
*N_sample_* = {30, 50, *and* 70}

With each combination of simulation parameters, we ran the simulation 20 times to obtain confidence intervals. DReAmiR was run using default parameters. As a comparison, we also evaluated the performance of two other methods using the same simulated dataset: (1) comparing the correlation density between groups using ANOVA, and (2) and summarizing the p-values with Edgington’s method^30^ implemented in the metap R package^31^ from two-group KS-test comparisons. All tests were performed with a significance value of p-value < 0.05. The performance metrics between methods were comapared using two-group Student’s T-tests.

### Cell-type-specific mmu-miR-155 KO RNA-seq data

The gene count tables for primary dendritic cells, B cells, CD4+ T cells, and macrophages from C56BL/6J wild-type and miR-155 KO mice were obtained from GSE116348^22^. The raw gene count data was normalized first using the Trimmed Mean of the M-values and transformed into log10 counts per million (logCPM)^32,33^. Then, genes with mean expression across samples equal to or less than 1 and interquartile range (IQR) equal to 0 were removed. The filtered count table with 10843 genes was TMM normalized again before conducting differential expression analysis.

Differential expression analysis was conducted within each cell-type separately. Within samples from each cell-type, the normalized expression data were first voom transformed^34^. Gene association with KO treatment was calculated by comparing the samples from the mmu-miR-155 KO group to the WT sample group using limma R package^33^. Genes were ranked by their association with mmu-miR-155 KO using t-statistics to generate cell-type specific rank lists. Signature genes were selected using log fold-change < 0 and FDR <= 0.05. TargetScan Mouse v7.1^13^ was used to predict the target genes for mmu-miR-155-5p (the major mature miRNA from pre-miR-155). To perform the max-dES target prioritization for each cell-type, gene rank lists generated from the other three cell-types together were used as the reference. Functional enrichment analysis of the mmu-miR-155 target genes was conducted using the hypeR R package^35^.

### Breast cancer molecular subtype analysis

miRNA and gene expression data and breast cancer molecular subtypes for TCGA BRCA samples were obtained using TCGAbiolinks^36^. We kept samples with both the miRNA and gene expression data from tumor samples and with PAM50 molecular subtypes (N=1064). miRNA and gene filtering was conducted using the same method described above, yielding 13846 genes and 584 miRNAs. Additionally, residual expression levels were calculated adjusting for the plate number using edgeR R package^32^, which were used for constructing correlation matrices per molecular subtypes. miRNA predicted target genes were queried from TargetScan v7.2^13^ and only the conserved target sites were used.

RNA-seq profiles of hsa-miR-23b perturbation assays (either hsa-miR-23b over-expression or hsa-miR-23b sponge vector transfection) and controls in MCF-7 and MDA-MB-231 cell lines were obtained from GSE37918^37^. The rank lists were generated using fold-change in each cell-line by comparing to the corresponding controls and signs were set that the negative association indicated a gene being regulated by miR-23b.

### Data and Code Availability

All data used in this project were publicly available. DReAmiR is an R package and can be downloaded from https://github.com/ningb/DReAmiR.

## Results

### DReAmiR method overview

A toy example is described in **Figure 1** to outline the basic workflow of the DReAmiR package. With the user-provided miRNA and gene expression matrix and the group label, DReAmiR computes the miRNA-gene correlation matrices for each group. DReAmiR also generates the corresponding miRNA-target gene matrix for the miRNAs and genes based on the intersection of miRNA target databases that the user specifies. SpQN can be performed as an optional step to adjust the correlation estimations and remove the mean-correlation relationship (**Figure 1a**). Next, DReAmiR performs differential regulation analysis for each miRNA using AD test (**Figure 1b**). By default, the genes are ranked by their correlation coefficients with miRNA within each group based on the correlation matrices generated in the previous step. Alternatively, the user can construct the gene rank list manually using other metrics such t-statistic from perturbation experiments. By comparing the negative enrichment pattern of target genes along the gene rank list by their correlation with the miRNA across groups rather than simply comparing the correlation densities, DReAmiR better captures the miRNA rewiring events and is less affected by the sample sizes. Parallel computation is implemented to speed up the process when multiple cores are available. An enrichment plot can be plotted to help visualize the difference in the enrichment patterns of target genes between groups. In the example shown in **Figure 1b**, the target genes of this miRNA were strongly negatively enriched in group A and B, but less in group C.

After getting a list of miRNA with significant context-specific targets, it might be important to identify group specific target genes for each miRNA. DReAmiR provides two independent methods to address this target prioritization step (**Figure 1c**). The graph embedding and node clustering method focuses on all predicted target genes in the leading-edge and aims to label each target as specifically regulated in one group, shared by multiple groups, or common across all. In the given example, four clusters of target genes were identified for the miRNA, where three of them were group-specific and one was shared across all groups. The max-dES method focuses on one particular group and aims to find the set of predicted target genes that generate the largest difference between this and the reference group. Assuming the group A was the group of interests, we combined the group B and C to generate the background rank list. While the resulted group-specific target genes from two methods could be overlapped, the user can choose which method to use depending on the specific biological question and study design.

### Benchmark on simulation data and comparison vs. other methods

Biological network algorithms are typically tested using ground truth data where the edges between individual nodes are known and experimentally validated. Yet, no such dataset is available for miRNA differential regulation and a method for similar purposes has not been developed to the best of our knowledge. Thus, we aimed to evaluate DReAmiR based on its robustness and the biological plausibility of its results.

To evaluate the ability of DReAmiR to identify miRNAs with context-specific target genes across multiple groups, we simulated gene and miRNA expression data with both true positive and true negative miRNAs and predicted target gene information (**Figure 2a**). Then, we performed DReAmiR with default settings 20 times for each parameter combination. DReAmiR demonstrated good performance under most scenarios, and the F1 score and TPR increased with *N_sample_* and *N_target_* (**Figure 2b** and **Supplementary Fig1**). Among these three parameters, the TPR and F1 score of DReAmiR performance were most strongly affected by the *P_variable%_*. Based on observations from Hsin JP *et al*.^22^, the majority of predicted targets are group-specific, suggesting DReAmiR should perform well in realistic settings, especially when *N_sample_* and *N_target_* are large enough. In the meantime, the FPR remains low (most less than 5%) regardless of the simulation parameters tested.

In addition, we also evaluated the performance of two alternative strategies in detecting miRNA regulatory rewiring across multiple groups, using the same simulated dataset. First, we used ANOVA to compare the correlation density distribution of predicted target genes for each miRNA, representing the underlying model of general gene-gene network-based algorithms, such as DGCA^7^. Second, we used the pair-wise group comparison function in DReAmiR based on KS-test and summarized the p-values using Edgington’s method, mimicking methods that only allow two-group comparison. Under all simulated parameters, DReAmiR outperformed these two methods (**Figure 2b** and **Supplementary Fig 1**). More speicifically, the F1 score and TPR from DReAmiR were significantly higher than those from ANOVA or p-value summation when *P_varible%_* or *N_target_* is large (**Supplementary Table 1**). Notably, the FPR from the ANOVA method is generally significantly higher than from DReAmiR. In contrast, while the p-value summation method can achieve relatively low FPR, the F1 score and TPR are much lower than DReAmiR. Similar difference in performance between the three methods was observed, where the sample size was different across groups (**Supplementary Figure 2**), suggesting that DReAmiR is not strongly affected by the sample size bias in correlation estimation. The simulation results suggested that DReAmiR is better at identifying miRNA with context-specific target genes than other methods and can achieve good performance under reasonable conditions (sample size, target number, and effect size) while maintaining high specificity.

### DReAmiR identified mmu-miR-155 cell-type-specific functional pathways

Having benchmarked DReAmiR in simulated data, we next sought to evaluate whether DReAmiR can identify miRNA known to regulate different targets in different contexts. We picked mmu-miR-155 as an example since the rewiring of the mmu-miR-155 regulatory network across immune cell types has been clearly demonstrated, and its functions in immune cells have been extensively studied^22,38,39^. Using all mmu-miR-155 predicted target genes as the target set (N=430) and the average logFC from differential expression analysis (comparing samples in the WT to the KO group) for generating the rank list per cell-type, differential regulation analysis was performed and the rank list from each cell-type was shuffled 500 times to calculate the permutation p-value. mmu-miR-155 regulated different target genes across four immune cell-types (**Figure 3b**; Permutation p-value = 0.01). In contrast, the average logFC distribution of mmu-miR-155 predicted target genes were not significantly different between four immune cell-types (**Figure 3a**; ANOVA p-value=0.08). These observations suggest DReAmiR may discover miRNAs with context-specific targets that are not captured by comparing average association strength.

We then used the max-dES method to prioritize group-specific target genes and examine whether the prioritized target genes were associated with cell-type-specific functions. The max-dES yielded 55, 40, 43, and 58 predicted target genes for B-cell, dendritic cell, macrophage, and CD4 T-cell, respectively, when each cell-type was compared to the other three as reference (**Figure 3c**). Target gene expression per cell-type was not strongly associated with whether a target gene was prioritized for a cell-type (**Supplementary Fig 3**) as previously suggested^22^, indicating the cell-type specific regulatory behavior of miR-155 was not solely determined by the target gene expression level. Notably, the rankings of the prioritized target genes overlapped between the group of interests and the reference group, meaning max-dES did not simply prioritize the top-ranking target genes. Functional pathway enrichment analysis was then performed on the prioritized target genes and the known pathways related to mmu-miR-155 cell-type-specific functions were among the top significantly enriched pathways (**Figure 3d**). For example, proliferative and apoptotic signaling pathways in B cells^40,41^, cytokine and co-stimulation related pathways in dendritic cells^42,43^, and activation and differentiation pathways in CD4+ T cells^44,45^. However, such enrichment was much weaker for the differentially expressed (logFC < 0 and FDR <= 0.05) miR-155 target genes (N=36, 110, 6 and 13). It is also worth noting that while it may be possible to alter the threshold for differential expression analysis to achieve similar functional enrichment results, no parameter tuning was needed for DReAmiR to generate such results. Taken together, this evidence suggested that DReAmiR can better identify miRNA rewiring events and the targets prioritized from DReAmiR are biologically informative.

### DReAmiR identified BRCA subtype-specific targets from bulk RNA-seq data

We next sought to demonstrate DReAmiR in a realistic use case, where researchers want to identify miRNAs with significant context-specific targets from sample-matched bulk miRNA and mRNA expression profiles between cancer subtypes, and validate the candidates through *in vitro* experiments. We performed the differential regulation analysis in TCGA BRCA data between five PAM50 breast cancer subtypes^46,47^, which yielded 10 miRNAs with significant context-specific target genes (**Figure 4a**; Permutation FDR <= 0.1). The mean expression levels of these miRNAs within each subtype showed very different patterns compared to their ES values, suggesting context-specific target gene regulation is not directly driven by differential expression levels. Then, we prioritized subtype-specific target genes for each molecular subtype using the graph embedding and node embedding method. The expression correlation densities between miRNA and the prioritized target genes for each subtype were significantly lower among the samples of each subtype compared to all other subtypes (**Figure 4b**; one-tail KS test, p-values < 0.01), suggesting the target genes prioritized by DReAmiR were specific to each subtype.

**Fig 4.**
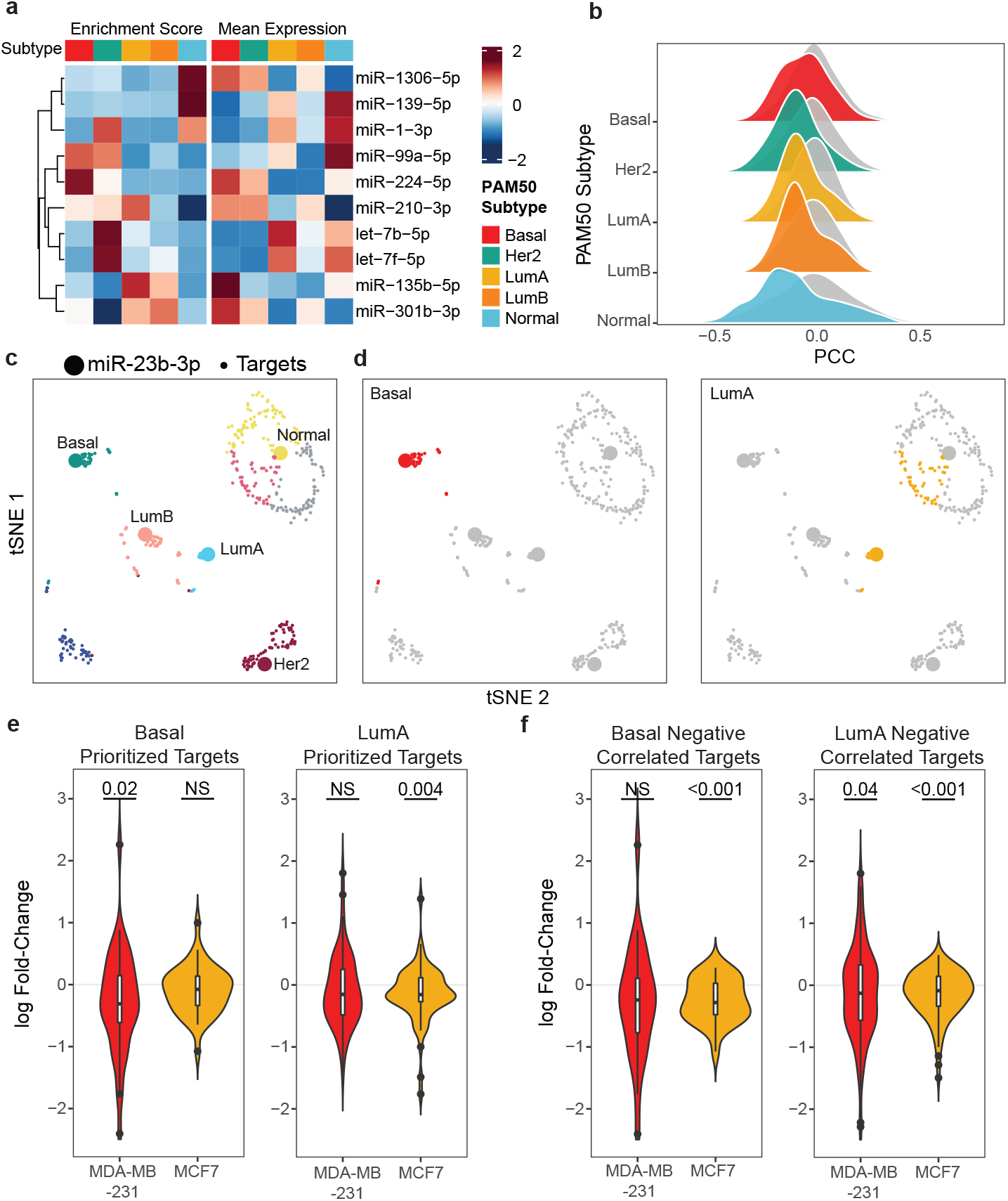
The miR-23b prioritized targets from TCGA BRCA are uniquely altered by perturbation in the cell line of the same subtype. **a.** Heatmap of the ES and mean expression levels of differentially regulating miRNAs in TCGA BRCA data by the PAM50 subtypes. The rows were scaled and clustered by the ES values. **b.** Correlation densities between significantly differentially regulating miRNAs and their prioritized target genes for each subtype (colored) or the prioritized target genes for the other four subtypes (grey). Comparisons within all subtypes were significant (one-tail KS-test p-value < 0.05). **c-d.** tSNE plots were generated using 10 embedded features for the leading-edge genes from hsa-miR-23b differential regulation analysis. Points were colored by fuzzy k-means clusters (**c**) and subtype assignment to either basal or luminal A subtype (**d**). The number of genes assigned to Basal subtype=42, Her2=128, LumA=72, LumB=64, Normal=261. The size of the points depicted the node type. **e.** LogFC densities of hsa-miR-23b prioritized target genes in MCF7 and MDA-MB-231 perturbation assays. **f.** LogFC densities of predicted target genes that were significantly negatively correlated with hsa-miR-23b in TCGA BRCA RNA-seq data in MCF7 and MDA-MB-231 perturbation assays.

To support that the prioritized target genes are indeed altered in a subtype-specific fashion, we examined whether the target genes prioritized for a breast cancer subtype were specifically altered with miRNA expression manipulation in the cell line of the same subtype. hsa-miR-23b was taken as an example since it is the only miRNA that we found perturbation assay with transcriptomic profiles^37^ in cell lines of different breast cancer subtypes: MCF-7 for the luminal A subtype and MDA-MB-231 for the basal subtype^48^, even though hsa-miR-23b does not appear to regulate different targets in the different breast cancer subtypes in the TCGA BRCA data (permutation FDR = 0.48).

Fuzzy k-means clustering on the graph embedding features derived from the TCGA BRCA data revealed eight different clusters. hsa-miR-23b in the five breast cancer subtypes were in separate clusters (**Figure 4c**). One cluster (42 predicted target genes) was labeled as unique to the basal subtype, and two clusters (72 predicted target genes) were labeled to be associated with Luminal A subtype with one shared with the normal subtype (**Figure 4d**). Genes in the basal subtype cluster were enriched in oxidative phosphorylation and Kit signaling pathways, and those in luminal A subtype clusters were enriched in citrate cycle and interleukin signaling pathways (**Supplementary Table 2**; hypergeometric test p-value < 0.01).

We next tested whether the targets of hsa-miR-23b that are differentially expressed in Basal and LumA breast cancer in the TCGA BRCA data show cell-line subtype-specific response to hsa-miR-23b overexpression *in vitro*. In the *in vitro* hsa-miR-23b overexpression experiment, the logFC of the basal-specific cluster genes were significantly lower than zero in MDA-MB-231, but not MCF7, suggesting these genes were specifically suppressed by hsa-miR-23b among the samples of the basal subtype (**Figure 4e**; one-tail t-test p-value < 0.05). A similar observation was seen for the luminal A specific gene cluster which were repressed by hsa-miR-23b overexpression in MCF7, but not MDA-MB31 (**Figure 4e**; one-tail t-test p-value < 0.05). In contrast, the logFC densities of the predicted target genes that were significantly negatively correlated with hsa-miR-23b (N=27) in the BRCA luminal A subtype samples were significantly lower than zero in both MDA-MB-231 and MCF-7 (**Figure 4f**; one-tail t-test, p-value < 0.05), while significant change was seen for the negatively correlated hsa-miR-23b predicted target genes in the BRCA basal subtypes only in MCF7 (one-tail t-test, p-value < 0.01) but not in MDA-MB-231. These observations highlighted the utility of DReAmiR for identifying cancer subtype-specific miRNA target genes and selecting uniquely regulated miRNA target genes from bulk expression data for *in vitro* studies.

## Discussion

Understanding miRNA regulatory network differences between cell types or disease states is essential to investigate the activities of miRNAs and their regulated gene programs. Existing methods were mostly developed for gene-gene networks, and fail to incorporate the predicted target information and the gene expression suppression nature of miRNAs. Furthermore, these methods often end at the rewiring detection step and do not quantitatively identify the group-specific targets of the regulators. In order to address these shortcomings, we created DReAmiR, a novel computational tool for characterizing miRNA-mediated regulatory network rewiring. First, DReAmiR calculates the miRNA-gene correlation matrix for each group and corrects the bias from the mean-correlation relationship using SpQN. Then, utilizing a GSEA-like model, DReAmiR detects miRNA with group-specific target regulation across multiple groups by comparing the negative enrichment patterns of the predicted target genes within gene rank list based on miRNA-gene correlation coefficients per group. Finally, DReAmiR identifies group-specific target genes through target prioritization.

Correlation estimation between miRNA and gene bulk expression profiles are affected by various technical factors, including expression levels and sample size. The accuracy of the differential regulation analysis depends on the ability to estimate miRNA-gene expression correlation free from such potential bias. Mean-correlation relationship was first described by Wang Y *et al*.^25^, and described a bias from the noise introduced during the high-throughput sequencing process that makes the absolute correlation coefficients estimated between highly expressed genes appear to be larger than those more lowly expressed. DReAmiR adopts and modifes the Wang Y *et al*. method to remove such bias before running the differential regulation analysis, such that the expression level of miRNA and gene may have less effect on downstream analysis.

Through testing our method in simulated data, we showed DReAmiR can achieve high performance under various conditions when correlation is used for ranking genes, particularly with a large number of predicted target genes and larger sample size, while maintaining low FPR. Compared with other strategies for modeling miRNA rewiring events, such as comparing correlation densities or summing p-values from two-group comparisons, the differential regulation analysis achieved better performance across different parameters settings, including various effect strength and sample sizes.

Using expression and mRNA-binding profiles from mmu-miR-155 KO mice, Hsin P *et al*. showed that the regulatory network rewiring of mmu-miR-155 between immune cell-types is a result of switching of binding targets^22^. Furthermore, while all the predicted target genes are being suppressed to some degree, the cell-type-specific targets of mmu-miR-155 have a stronger association with mmu-miR-155 KO in that cell-type compared to all predicted targets. Hence, simply comparing the association strength, either based on fold-change in perturbation assays or correlation coefficient, between a miRNA and all of its predicted targets lack the sensitivity for detecting miRNA-related rewiring events. To better identify miRNAs with context-specific target genes, DReAmiR uses a framework similar to gene-set enrichment analysis (GSEA)^49^. One major task for GSEA is to examine whether genes of interest (the gene set) are concordantly positively or negatively enriched along a gene rank list, where the genes are ranked by their association with certain phenotypes. In DReAmiR, we use an Anderson Darling’s test and ask whether the predicted target genes of a miRNA (a target set) are similarly negatively enriched along multiple rank lists where genes are ranked by their association to a miRNA in each group. Applying DReAmiR to the RNA-seq data from Hsin P *et al*, we identified mmu-miR-155 as an miRNA with significant context-specific targets. Yet, simply comparing logFC densities of predicted target genes failed to detect the rewiring event. These results demonstrate that DReAmiR, by modeling the negative enrichment pattern of miRNA’s predicted target genes in the correlation rank list across multiple groups, is usable and suitable for detecting miRNA regulatory network rewiring.

A typical task for studying miRNA is to identify top candidate target genes *in silico* and validate the findings through *in vitro* or *in vivo* assays. We demonstrated the utility of DReAmiR for such real-world scenario with the TCGA BRCA subtype analysis. Conventionally, putative target genes are identified by setting a significance threshold to filter a set of genes through subtype-specific negative correlation analysis in bulk expression profiles. However, such a task may often fail because it does not inform whether the targets identified are specific to a group, and setting a hard threshold introduces unnecessary bias. Indeed, the target genes identified using this strategy did not show subtype-specific expression changes in cell lines with miR-23b perturbations. In contrast, DReAmiR does not set a specific hard threshold on the correlation coefficient nor does it significance level to filter the target genes. More importantly, the miRNA target genes identified through target prioritization for each cancer subtype were altered in a cancer subtype-specific fashion in cell line perturbation assays and were enriched for different functional pathways. Notably, hsa-miR-23b regulation over several of the prioritized target genes, including PIL3R3 and PAK2, have been suggested in breast cancer settings^37,50^. Yet, the BRCA subtype specific regulation was not extensively studied and may inform future investigations. These observations, which were not found by traditional methods, highlighted the unique advantage of DReAmiR.

DReAmiR offers two target prioritization methods to identify putative group-specific target genes for miRNAs: graph embedding and dES maximization. These two methods aim to answer similar questions but from different perspectives. Graph embedding aims to characterize the distance between the gene nodes and group-specific miRNA nodes in the embedded space, and assign target genes to one or multiple groups based on cluster membership probabilities. Thus, it globally examines all target genes and all groups at once. This is useful when users are interested in all experimental groups or disease subtypes, or targets shared between groups, as we showed in the TCGA BRCA subtype analysis. dES maximization, on the other hand, tries to iteratively search for a set of target whose normalized enrichment score along the rank list of the group of interests most different from the background. The background can either be another group that users want to make contrast with, or all other samples. This can be used when samples from multiple groups are similar based on prior belief and the analysis is logical as a two-group comparison. For example, in the mice immune cell types analysis, we identified the target genes strongly associated with mmu-miR-155 KO in one cell type of interests, with samples from the remaining three closely related cell types combined as the background. This should be used when there is one group that is particular interested to the users, such as a cancer-subtype with significantly worse prognosis comparing to others. While two methods often yield very similar results, we advise users to choose the method better suited to their experimental design and biological questions for the best interpretability.

Although DReAmiR was primarily developed for miRNA analysis, it can be extended for other types of transcriptomic regulatory networks and groups other than cancer subtypes or cell types, as long as a target set and group-specific rank lists could be defined. Also, we included a user-defined option to choose the direction of interest for enrichment such that either suppressive or activating regulators could be modeled during the target prioritization step. For example, DReAmiR can be used to examine the group-specific activities of RNA-binding proteins, long non-coding RNAs, or even TFs between multiple cancer subtypes. It can also be applied to study chromatin regulator behavior or histone modifications, where the chromatin-binding peak intensities are used for generating rank lists and peaks associated with certain target genes or functions as the target set. As an extension to gene set variation analysis^51^, DReAmiR can also answer a question like whether genes in a predefined gene set are similarly correlated with a phenotype across groups, similar to what we did for the mmu-miR-155 KO dataset.

There are several limitations for DReAmiR worth mentioning. First, DReAmiR does not provide a mechanism for why miRNA regulatory rewiring happens between conditions, and additional biochemical studies will need to be performed for this purpose. Also, even though we extensively validate DReAmiR results using known biological observations, there are limited examples of known context-specific miRNA regulation to serve as ground-truth for validating DReAmiR. With sequencing technology development, we hope methods such as cross-linking ligation and sequencing of hybrids^52^ (CLASH) can be improved such that the exact binding between all miRNAs and target genes could be examined for large sample size and single-cell level for DReAmiR validation.

In conclusion, DReAmiR is a new approach to characterize miRNA-mediated gene regulatory network rewiring across multiple groups from transcriptomic profiles. The method may offer novel insights into cell-type and cancer subtype-specific miRNA regulatory roles. In the future, we plan to apply the method to the premalignant bronchial lesion biopsy data and examine whether miRNA with differential regulatory roles may be associated with the molecular subtypes.

---

**Supplementary Fig 1.**
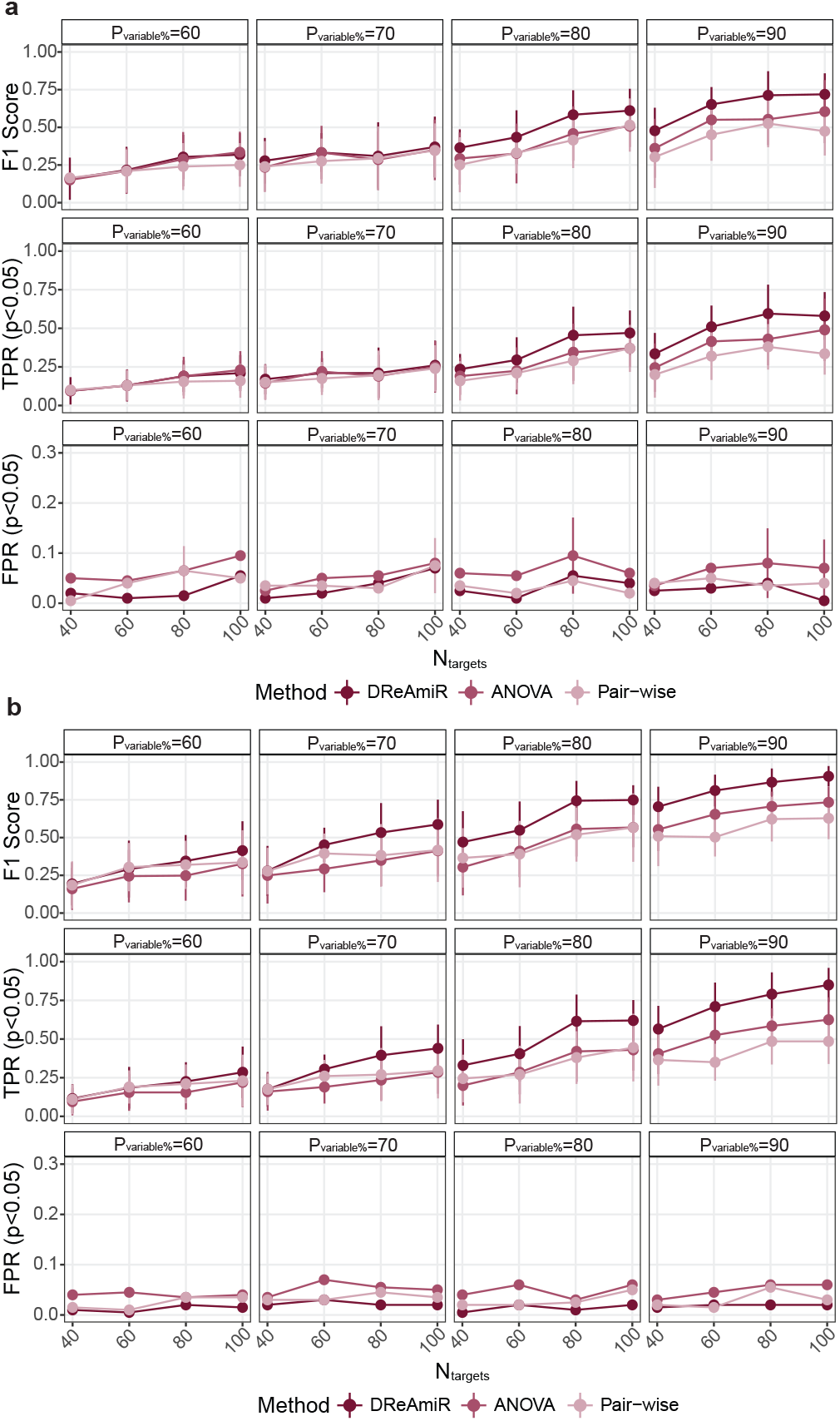
DReAmiR performance in simulated data with *N_sample_* equaled 30 (**a**) and 50 (**b**).

**Supplementary Fig 2.**
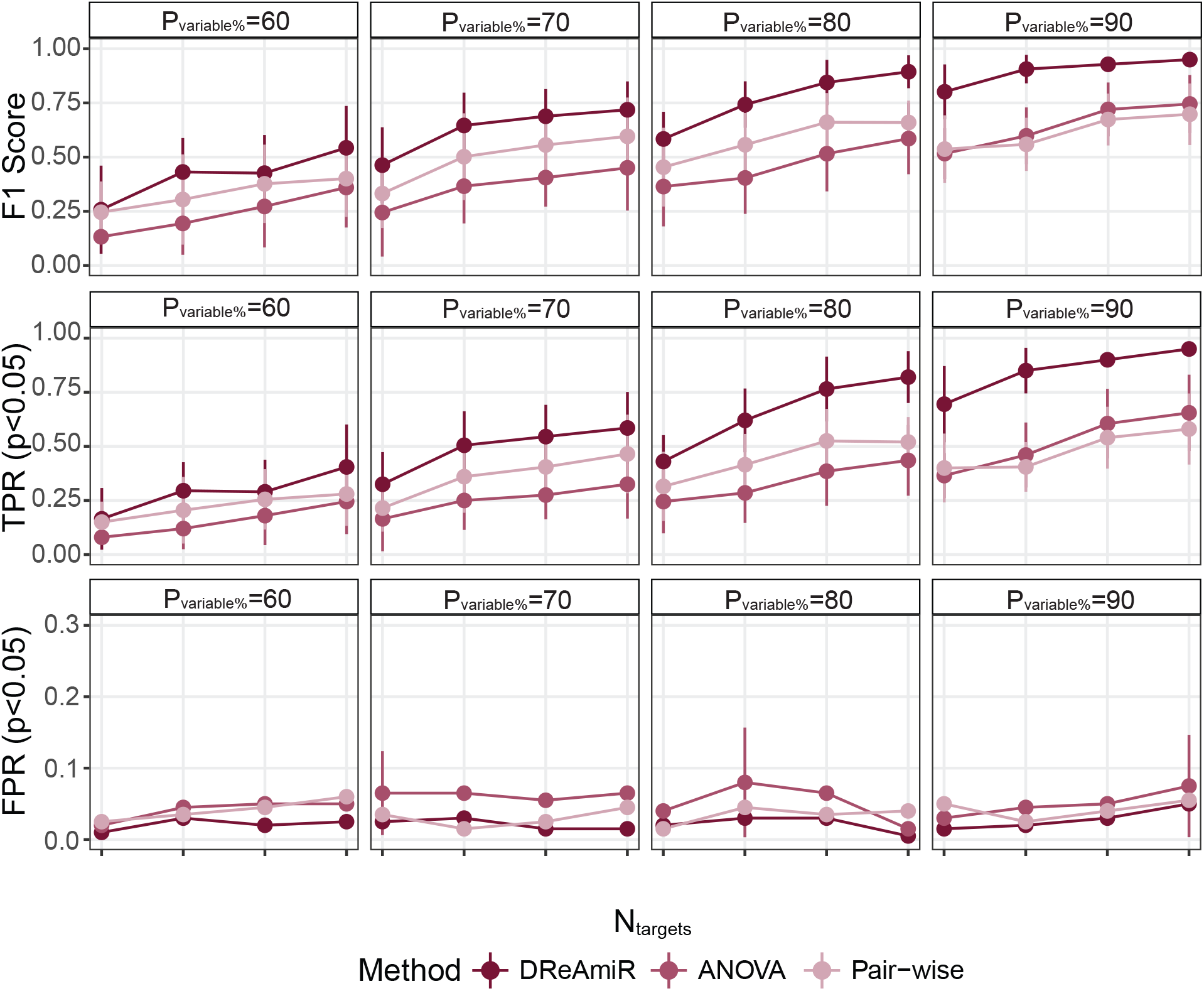
DReAmiR performance in simulated data with different sample sizes across groups (*N_sample_* =50, 60 and 70).

**Supplementary Fig 3.**
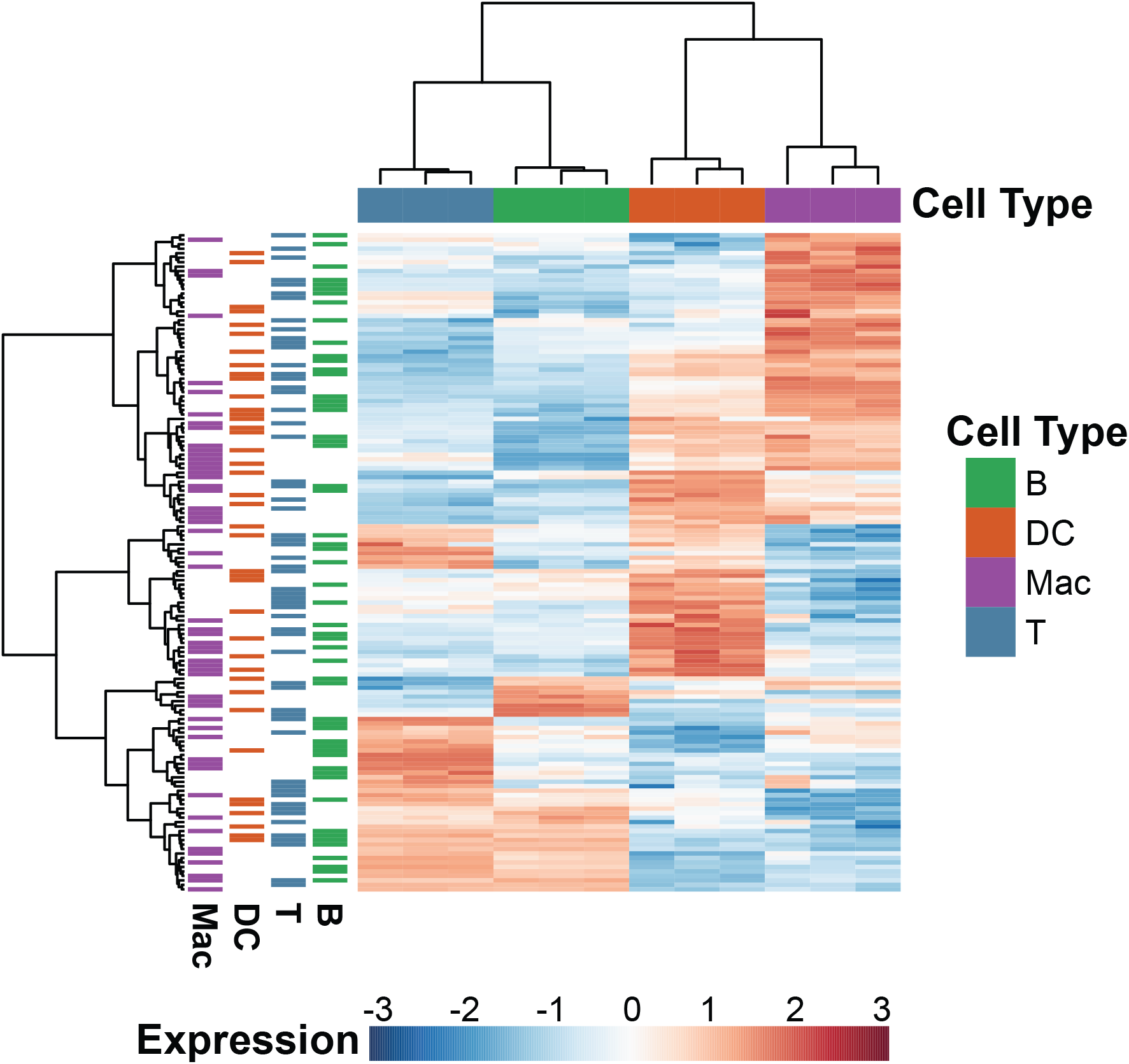
The expression level of miR-155 prioritized target genes across four mice immune cell-types of the control group. The color bar on the top showed the cell-types from the mice of the control group, and the color bar on the left of the heatmap showed which cell-type the prioritiozed target gene belonged to. The expression values were scaled per gene.

## References

1. Gerstein, M. B. et al. Architecture of the human regulatory network derived from ENCODE data. Nat. 2012 4897414 489, 91–100 (2012).

2. Lee, T. I. et al. Transcriptional regulatory networks in Saccharomyces cerevisiae. Science (80-.). 298, 799–804 (2002).

3. Aibar, S. et al. SCENIC: Single-cell regulatory network inference and clustering. Nat. Methods 14, 1083–1086 (2017).

4. Ding, K. F. et al. Network rewiring in cancer: Applications to melanoma cell lines and the cancer genome atlas patients. Front. Genet. 9, 228 (2018).

5. Assi, S. A. et al. Subtype-specific regulatory network rewiring in acute myeloid leukemia. 51, 151–162 (2019).

6. Bhardwaj, N., Kim, P. M. & Gerstein, M. B. Rewiring of transcriptional regulatory networks: hierarchy, rather than connectivity, better reflects the importance of regulators. Sci. Signal. 3, (2010).

7. Mckenzie, A. T., Katsyv, I., Song, W. M., Wang, M. & Zhang, B. DGCA : A comprehensive R package for Differential Gene Correlation Analysis. BMC Syst. Biol. 10, 1–25 (2016).

8. Zhang, J. et al. DiNeR: a Differential graphical model for analysis of co-regulation Network Rewiring. BMC Bioinformatics 21, 281 (2020).

9. Lou, S. et al. TopicNet: a framework for measuring transcriptional regulatory network change. Bioinformatics 36, i474–i481 (2020).

10. Tesson, B. M., Breitling, R. & Jansen, R. C. DiffCoEx: a simple and sensitive method to find differentially coexpressed gene modules. 11, 497 (2010).

11. Jonas, S. & Izaurralde, E. Towards a molecular understanding of microRNA-mediated gene silencing. Nature Reviews Genetics 16, 421–433 (2015).

12. Bartel, D. P. Metazoan MicroRNAs. Cell 173, 20–51 (2018).

13. Agarwal, V., Bell, G. W., Nam, J. & Bartel, D. P. Predicting effective microRNA target sites in mammalian mRNAs. 1–38 (2015). doi:10.7554/eLife.05005

14. Gebert, L. F. R. & MacRae, I. J. Regulation of microRNA function in animals. Nat. Rev. Mol. Cell Biol. 2018 201 20, 21–37 (2018).

15. DeVeale, B., Swindlehurst-Chan, J. & Blelloch, R. The roles of microRNAs in mouse development. Nat. Rev. Genet. 2021 225 22, 307–323 (2021).

16. Esquela-Kerscher, A. & Slack, F. J. Oncomirs — microRNAs with a role in cancer. Nat. Rev. Cancer 2006 64 6, 259–269 (2006).

17. Lin, S. & Gregory, R. I. MicroRNA biogenesis pathways in cancer. Nat. Rev. Cancer 15, 321–333 (2015).

18. Lionetti, M. et al. Identification of microRNA expression patterns and definition of a microRNA/mRNA regulatory network in distinct molecular groups of multiple myeloma. Blood 114, e20–e26 (2009).

19. Nam, J. W. et al. Global analyses of the effect of different cellular contexts on microRNA targeting. Mol. Cell 53, 1031–1043 (2014).

20. Lin, C. C. et al. Regulation rewiring analysis reveals mutual regulation between STAT1 and miR-155-5p in tumor immunosurveillance in seven major cancers. Sci. Reports 2015 51 5, 1–11 (2015).

21. Nowakowski, T. J. et al. Regulation of cell-type-specific transcriptomes by microRNA networks during human brain development. Nat. Neurosci. 21, 1784–1792 (2018).

22. Hsin, J. P., Lu, Y., Loeb, G. B., Leslie, C. S. & Rudensky, A. Y. The effect of cellular context on miR-155-mediated gene regulation in four major immune cell types. 19, 1137–1145 (2018).

23. Li, X. et al. High-Resolution In Vivo Identification of miRNA Targets by Halo-Enhanced Ago2 Pull-Down. Mol. Cell 79, 167–179.e11 (2020).

24. Helwak, A., Kudla, G., Dudnakova, T. & Tollervey, D. Mapping the Human miRNA Interactome by CLASH Reveals Frequent Noncanonical Binding. Cell 153, 654 (2013).

25. Wang, Y., Hicks, S. C. & Hansen, K. D. Co-expression analysis is biased by a mean-correlation relationship. bioRxiv 2020.02.13.944777 (2020). doi:10.1101/2020.02.13.944777

26. Tang, J. et al. LINE: Large-scale Information Network Embedding. WWW 2015 - Proc. 24th Int. Conf. World Wide Web 1067–1077 (2015). doi:10.1145/2736277.2741093

27. Ru, Y. et al. The multiMiR R package and database: integration of microRNA–target interactions along with their disease and drug associations. Nucleic Acids Res. 42, e133–e133 (2014).

28. Geraci, M. Linear Quantile Mixed Models: The lqmm Package for Laplace Quantile Regression. J. Stat. Softw. 57, 1–29 (2014).

29. Wenables, W. & Ripley, B. Modern Applied Statistics with S, Fourth edition. (Springer, New York, 2002).

30. Edgington, E. S. An additive method for combining probability values from independent experiments. J. Psychol. Interdiscip. Appl. 80, 351–363 (1972).

31. Dewey, M. Meta-Analysis of Significance Values [R package metap version 1.6]. (2021).

32. McCarthy, D. J., Chen, Y. & Smyth, G. K. Differential expression analysis of multifactor RNA-Seq experiments with respect to biological variation. Nucleic Acids Res. 40, 4288–4297 (2012).

33. Ritchie, M. E. et al. limma powers differential expression analyses for RNA-sequencing and microarray studies. 43, (2015).

34. Law, C. W., Chen, Y., Shi, W. & Smyth, G. K. Voom: Precision weights unlock linear model analysis tools for RNA-seq read counts. Genome Biol. 15, 1–17 (2014).

35. Federico, A. & Monti, S. hypeR: an R package for geneset enrichment workflows. Bioinformatics 36, 1307–1308 (2020).

36. Colaprico, A. et al. TCGAbiolinks: an R/Bioconductor package for integrative analysis of TCGA data. Nucleic Acids Res. 44, e71–e71 (2016).

37. Pellegrino, L. et al. miR-23b regulates cytoskeletal remodeling, motility and metastasis by directly targeting multiple transcripts. Nucleic Acids Res. 41, 5400–5412 (2013).

38. Mehta, A. & Baltimore, D. MicroRNAs as regulatory elements in immune system logic. Nat. Rev. Immunol. 16, 279–294 (2016).

39. Alivernini, S. et al. MicroRNA-155-at the critical interface of innate and adaptive immunity in arthritis. Front. Immunol. 8, 1932 (2018).

40. Zhu, F. Q. et al. MicroRNA-155 Downregulation Promotes Cell Cycle Arrest and Apoptosis in Diffuse Large B-Cell Lymphoma. Oncol. Res. 24, 415–427 (2016).

41. Yu, H. et al. MicroRNA-155 regulates the proliferation, cell cycle, apoptosis and migration of colon cancer cells and targets CBL. Exp. Ther. Med. 14, 4053 (2017).

42. Hodge, J. et al. Overexpression of microRNA-155 enhances the efficacy of dendritic cell vaccine against breast cancer. Oncoimmunology 9, (2020).

43. Lind, E. F. et al. miR-155 Upregulation in Dendritic Cells Is Sufficient To Break Tolerance In Vivo by Negatively Regulating SHIP1. J. Immunol. 195, 4632–4640 (2015).

44. Goncalves-Alves, E. et al. MicroRNA-155 controls T helper cell activation during viral infection. Front. Immunol. 10, 1367 (2019).

45. Chen, L., Gao, D., Shao, Z., Zheng, Q. & Yu, Q. miR-155 indicates the fate of CD4 + T cells. Immunol. Lett. 224, 40–49 (2020).

46. Bernard, P. S. et al. Supervised risk predictor of breast cancer based on intrinsic subtypes. J. Clin. Oncol. 27, 1160–1167 (2009).

47. Berger, A. C. et al. A Comprehensive Pan-Cancer Molecular Study of Gynecologic and Breast Cancers. Cancer Cell 33, 690–705.e9 (2018).

48. Dai, X., Cheng, H., Bai, Z. & Li, J. Breast Cancer Cell Line Classification and Its Relevance with Breast Tumor Subtyping. J. Cancer 8, 3131 (2017).

49. Subramanian, A. et al. Gene set enrichment analysis: a knowledge-based approach for interpreting genome-wide expression profiles. Proc. Natl. Acad. Sci. U. S. A. 102, 15545–50 (2005).

50. Zhang, H. et al. Genome-wide functional screening of miR-23b as a pleiotropic modulator suppressing cancer metastasis. Nat. Commun. 2, (2011).

51. Hänzelmann, S., Castelo, R. & Guinney, J. GSVA: Gene set variation analysis for microarray and RNA-Seq data. BMC Bioinformatics 14, (2013).

52. Helwak, A. & Tollervey, D. Mapping the miRNA interactome by cross-linking ligation and sequencing of hybrids (CLASH). Nat. Protoc. 2014 93 9, 711–728 (2014).

